# 3,3’-Diindolylmethane induces apoptosis and autophagy in fission yeast

**DOI:** 10.1101/2021.08.05.455326

**Authors:** Parvaneh Emami, Masaru Ueno

**Author notes:** Corresponding author: (MU).

## Abstract

3,3’-Diindolylmethane (DIM) is a compound derived from the digestion of indole-3-carbinol, found in the broccoli family. It induces apoptosis and autophagy in some types of human cancer. DIM extends lifespan in the fission yeast *Schizosaccharomyces pombe*. The mechanisms by which DIM induces apoptosis and autophagy in humans and expands lifespan in fission yeasts are not fully understood. Here, we show that DIM induces apoptosis and autophagy in log-phase cells, which is dose-dependent in fission yeast. A high concentration of DIM disrupted the nuclear envelope (NE) structure and induced chromosome condensation at an early time point. In contrast, a low concentration of DIM induced autophagy but did not disrupt NE structure. The mutant defective in autophagy was more sensitive to a low concentration of DIM, demonstrating that the autophagic pathway contributes to the survival of cells against DIM. Moreover, our results showed that the *lem2* mutant is more sensitive to DIM. NE in the *lem2* mutant was disrupted even at the low concentration of DIM. Our results demonstrate that the autophagic pathway and NE integrity are important to maintain viability in the presence of a low concentration of DIM. The mechanism of apoptosis and autophagy induction by DIM might be conserved in fission yeast and humans. Further studies will contribute to the understanding of the mechanism of apoptosis and autophagy by DIM in fission yeast and humans.

## Introduction

Apoptosis is a process that results in the death of damaged and unrepairable cells to maintain health in multicellular organisms [1]. It is a promising target for cancer therapy as it induces death in cancer cells [2]. Autophagy also recycles the cellular components; it can be induced by stress or nutrient starvation [3, 4]. Upregulation of autophagy extends the lifespan of the animal [5]. Autophagy induction can suppress the growth of cancer cells [6–9]; therefore, it has several potential benefits to human health, including longevity and cancer therapy.

3,3’-Diindolylmethane (DIM) is a compound derived from the digestion of indole-3-carbinol, found in the plants from the broccoli family, such as cabbage, broccoli, and rape [10]. Recent studies reported the anti-cancer effects of DIM through the induction of apoptotic cell death in breast cancer [11, 12], hepatoma [13, 14], prostate cancer [15, 16], or colon cancer [17–19]. DIM rapidly accumulates in the nuclear membranes (NM) of human breast carcinoma (MCF-7) cells after 0.5-2 hours [20]. However, the outcome of the accumulation of DIM in NM remains unclear. DIM induces autophagy in gastric cancer cells [9] and in ovarian cancer cells [6] through endoplasmic reticulum (ER) stress. However, it is not completely understood how DIM induces ER stress.

Fission yeast *Schizosaccharomyces pombe* is a powerful model organism to study apoptosis [21–24] and autophagy [25, 26]. Apoptosis is detectable in fission yeast by observing dead cells and nuclear fragmentation [24, 27, 28] that could be induced by different conditions. Overexpression of endogenous gene such as *calnexin* in fission yeast causes apoptotic cell death, which partially depends on the Ire1, the ER-stress transducer [28]. On the contrary, *dga1 plh1* double mutant, which is defective in lipid metabolic pathway, promotes apoptosis in stationary-phase [29, 30]. Also, some compounds such as terpinolene [27] and alpha-thujone [21] induce apoptosis in this organism.

Autophagy can be induced in fission yeasts [31, 32] via different conditions. Nitrogen starvation induces autophagy in fission yeasts, which generates a nitrogen source for adaptation [33, 34]. Sulfur starvation also induces autophagy in fission yeasts [35]. Autophagy contributes to maintaining cellular viability in both nitrogen [25] and sulfur starvation [35]. Atg8 is critical for autophagosome formation in fission yeasts [8, 26] and used as a general autophagy marker that is distributed in the cytoplasm but forms bright foci when autophagy is induced by nitrogen starvation [31, 34, 36].

Nuclear envelope (NE) consists of the inner and outer NM, and the nuclear pore complex (NPC) in fission yeast [37]. One of the inner NM proteins in fission yeast is Lem2 that contributes to the NE integrity [38], maintaining the boundary between NE and ER [39] and controlling NM flow [40]. In *lem2Δ* cells, the distribution of ER protein such as Ish1 is affected [39]. Also, Lem2 recruits Cmp7 for NE closure by ESCRT-III [41]. ESCRT-III or the endosomal sorting complex required for transport-III is responsible for sealing a membrane rupture not only during mitosis but also during interphase to maintain NE integrity in fission yeast [39, 41, 42], which needs Vps4 to complete the sealing [41, 43].

There is not enough information about the effects of DIM on fission yeasts yet, except that it increases chronological lifespan (CLS) [44]. Here, we show that DIM decreases fission yeast viability at an early time point. We describe DIM as an apoptosis and autophagy inducer in this organism. Our results suggest that NE could be one of the early targets of DIM. We hope that these results help future studies, especially for cancer-related research.

## Materials and methods

### Procurement of yeast strains and construction of *Htb1-GFP, GFP-atg8, ire1Δ, atg7Δ and lem2Δ* strains

Fission yeast strains used in our experiments are listed in Table 1.

**Table 1.** The strains used in this paper

| Name | Genotype | Source |
| --- | --- | --- |
| 975 | *h+* | M. Yanagida |
| FY7455 | *h+ leu1-32 ura4-D18 his7 lys1-131* | NBRP |
| *htb1-GFP* | *h+ leu1-32 ura4-D18 his7 lys1+::(hta1 GFP-htb1)* | This paper |
| FY17186 | *h90 ade6-216 leu1-32 lys1-131 ura4-D18* | NBRP |
| *GFP-atg8* | *h90 ade6-216 leu1-32 ura4-D18 lys1+Pnmt41-GFP-Atg8-1* | This paper |
| *GFP-atg8 ire1Δ* | *h90 ade6-216 leu1-32 ura4-D18 lys1+Pnmt41-GFP-Atg8-1 ire1.m6::Kanr* | This paper |
| *ire1Δ* | *h90 ade6-216 leu1-32 ura4-D18 lys1-131 ire1.m1::Kanr* | This paper |
| *atg7Δ* | *h90 ade6-216 leu1-32 ura4-D18 lys1-131 atg7-m1::Natr* | This paper |
| FY29360 | *h90 hat1+::GFP-Kanr ark1::Kanr-Prad21-ark1 + atg7-d1::Natr* | NBRP |
| yPK002 | *h+ ade6-M210 ura4-D18 leu1-32 ire1::Kanr* | P. Walter |
| YT2416 | *h- lem2Δ::Kanr* | Y. Hiraoka |
| 80-G10 | *h+ ade6-210 leu1-32 cut11::cut11+-GFP-HA-Kanr sid4+-mCherry<<Natr* | lab stock |
| 81-D02 | *h+ leu1-32 cut11::cut11+-GFP-HA-Kanr sid4+-mCherry<<Natr lem2Δ::Kanr* | This paper |
| 1-1-1 | *h+ ade6-210 leu1-32 lys1 ura4 sod2.prox::Kanr-ura4+-lacOp his7+::lacI-GFP sid4+-GFP-natBsd gar2+-mCherry-HphBsd* | Ito et al. 2019 |
| 78-A02 | *h+ ade6-210 leu1-32 lys1-131 ura4-D18 sod2.prox::Bsd-ura4+-lacOp his7+::lacI-GFP sid4+-GFP-natBsd gar2+-mCherry-hphMX6* | This paper |
| 78-D01 | *h+ ade6-210 leu1-32 lys1-131 ura4-D18 sod2.prox::Bsd-ura4+-lacOp his7+::lacI-GFP sid4+-GFP-natBsd gar2+-mCherry-HphBsd lem2::Kanr* | This paper |

Htb1-GFP strain was created by the introduction of NsiI partial-digested htb1GFPC plasmid, obtained from Dr. Matsuda and Dr. Hiraoka [45], to the FY7455 strain using *lys1*^+^ marker for selection.

To express GFP-tagged Atg8 in the wild-type strain (FY17186), we integrated EcoRI digested pHM43 plasmid containing a GFP-Atg8 gene with the *nmt41* promoter, gifted by Dr. Yamamoto [33], using *lys1*^+^ marker for selection.

To construct the *ire1Δ::Kanr* strain, we amplified the genomic DNA by polymerase chain reaction (PCR) using genomic DNA obtained from yPK002 as a template, a gift from Dr. Walter [46], and the primers 5’-TGGATGACTATACCCAAAGC-3’ and 5’-ATCCAACGATCCCACAAGCG-3’. The resulting PCR product was introduced into the FY17186 strain by using *S. pombe* Direct Transformation Kit Wako (FUJIFILM Wako, Osaka, Japan). The deletion was confirmed by a PCR.

A similar strategy was adopted to create the autophagy mutant *atg7Δ::Natr*, using the primers 5’-ATACGTAGAACTGCGGTGAG-3’ and 5’-CAAATGCAACTTCAGGATCC-3’ to amplify the mutated genomic DNA from *atg7Δ* strain (FY29360) and introduce it to the wild-type strain, FY17186. The deletion was confirmed by a PCR.

To construct *lem2Δ::Kanr* mutant for the viability assay, first, the strain named 78-A02 was created as a wild-type strain by replacing *Kanr* with *Bsdr* in the strain 1-1-1 shown in Ito et al. [47]. First, *Bsdr* fragment-containing plasmid was created by replacing the *Natr* gene in pNATZA13-mCherry-*atb2*^+^, a gift from Dr. Y. Watanabe and Dr. T. Sakuno [48] with *Bsdr* gene by ligation of SacI and BglII digested pNATZA13-mCherry-*atb2*^+^ with *Bsdr* gene, which was amplified from pSVEM-Bsdr as a template, a gift from Dr. A. Stewart [49], with the primers 5’-CCCCTCACAGACGCGTCACTCAACCCTATCTCGG-3’ and 5’-ATCCGCCGGTACGCGTCTCGAAATTAACCCTCAC-3’, using In-Fusion HD Cloning Kit (Takara bio, Shiga, Japan). In the next step, the DNA fragment containing the *Bsdr* gene was amplified using 5’-GACATGGAGGCCCAGAATAC-3’ and 5’-TGGATGGCGGCGTTAGTATC-3’. The resulting DNA fragment was used for transformation to replace *Kanr* with *Bsdr* in the strain 1-1-1. The resulting strain 78-A02 was used to delete with *lem2Δ::Kanr. lem2Δ::Kanr* DNA fragment was amplified using the primers: 5’-CCCTAATGATCATGGATTCTGT-3’ and 5’-ACTATGGATGCCTATTTTCCC-3’ using genomic DNA of the YT2416 strain, obtained from Dr. Hiraoka [50], as a template. Then, the PCR fragment was introduced to the strain 78-A02 to create strain 78-D01. To construct *cut11-GFP* strain with *lem2Δ:Kanr*, the lab stock strain (80-G10) was mated with the *lem2Δ:Kanr* strain YT2416, resulting in strain 81-D02.

### Viability assay

In viability assay, 3% glucose Yeast extract with adenine (YEA) liquid medium, and YEA plates were employed. A composition of 0.5% yeast extract, 3% glucose, and 40 mg/ml adenine was used for the liquid YEA medium or the YEA plate with 2% agar. For viability assay, the fission yeast cells were grown in an 8 ml liquid YEA medium overnight at 30°C (12 ~ 15 hours) to get the log-phase cells (0.5 ~ 1×10^7^ cells/ml) designated as day 0. As seen in Fig. 1A, the culture was divided into two flasks. The same volume of Dimethyl sulfoxide (DMSO) or 20 μg/ml DIM, purchased from Combi-Blocks (San Diego, USA) dissolved in DMSO were added to each flask. On day 1, which was 24 hours after incubation with the drugs, cell number was adjusted to 1 × 10^7^ cells/ml and used for viability assay with five-fold serial dilutions. The spotted plates were incubated at 30°C for 3 to 5 days to check cell viability. Every 24 hours, the viability assays were repeated.

**Figure 1.**
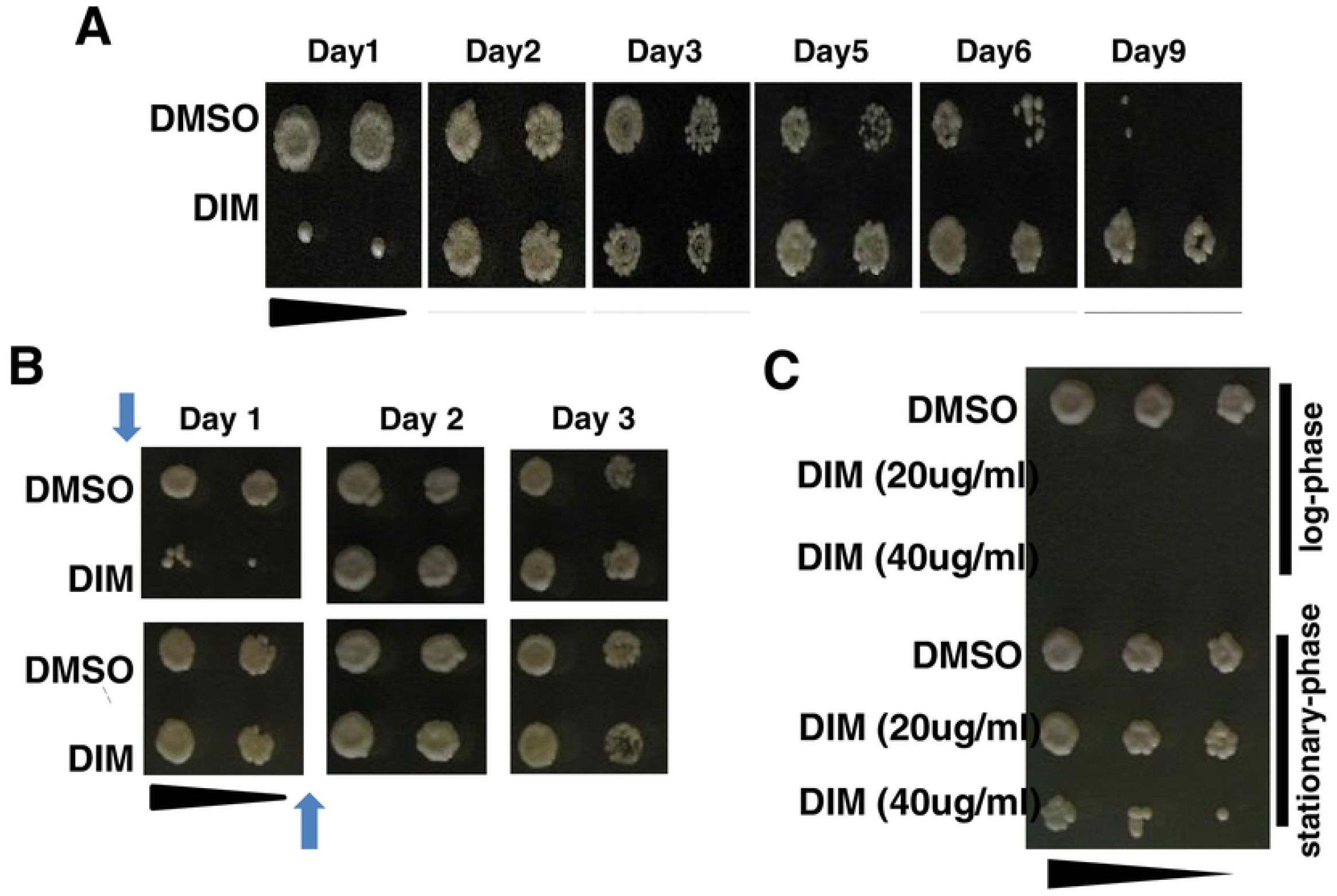
DIM inhibits the cell viability of log-phase cells. **(A)** 20 μg/ml 3,3’-Diindolylmethane (DIM) was added to the log-phase cells (~0.5-1 x 10^7^ cells/ml) and the viability was studied by spot assay with five-fold serial dilutions. Dimethyl sulfoxide (DMSO) was used for the control experiment. The same cell concentration was spotted on YEA plates to regularly check the cell viability till day 9. Day 1 means 24 hours incubation after adding the drug. **(B)** 20 μg/ml DIM was added to the stationary-phase cells and the viability was studied by a spot assay. Top panel: DIM was added to log-phase cells (0.5 ~ 1 x 10^7^ cells/ml) as a control (before day 1). Bottom panel: log-phase cells were cultured for 24 hours. Then, DIM was added to the stationary-phase cells (~1 x 10^8^ cells/ml). The same cell concentrations were spotted on YEA plates to regularly check the cell viability till day 3. Arrows show the time of adding DIM. **(C)** To perform the acute viability assay, the same concentration of log-phase cells and stationary-phase cells (1 x 10^7^ cells/ml) were incubated in the presence of DIM and DMSO for 10 minutes. Treated cultures were spotted and incubated at 30°C for 3 to 5 days to capture images. See materials and methods for details.

To compare the effects of DIM on stationary-phase cells with that on log-phase cells, day0 cells, shown in Fig. 1A, were cultured for 24 hours, which contained about ~1 × 10^8^ cells/ml designated as day 1. Then, DIM or DMSO were added (Fig. 1B). As a control, DIM or DMSO were added to the day0 culture. In viability assay, as seen in Fig. 1B, the cell number was adjusted to 1×10^7^ cells/ml and used for viability assays with five-fold serial dilutions. Every 24 hours, the viability assay was repeated. The viability assays, shown in Fig. 5 and Fig. 6, were performed as shown in Fig. 1A on fresh YEA plates containing 5 μg/ml DIM or 10 mM 2-mercaptoethanol (2-ME).

### Acute viability assay

The log-phase and stationary-phase cells were prepared as shown in Fig. 1A and 1B. The cell concentrations were adjusted to 1 × 10^7^ cells/ml. Immediately, DMSO or DIM (20 and 40 μg/ml) were added to the cultures and incubated at 30°C for 10 minutes. After washing the drugs, 100 μl of the same concentration (1×10^7^ cells/ml) of treated cultures were used for viability assay with five-time serial dilutions on YEA plates (Fig. 1C).

### Apoptosis detection by acridine orange/ethidium bromide (AO/EB) and 4’,6-diamidino-2-phenylindole (DAPI)

Dual staining by AO/EB was conducted to detect dead cells as previously described [27]. Briefly, after precipitation and washing the cell with PBS (pH: 7.4), cells were resuspended in 100 μl PBS with 5 μl of AO/EB mixture (AO 60 μg/ml: EB 100 μg/ml). Five minutes after incubation in the darkroom at room temperature, cells were washed twice with PBS and imaged by a Zeiss GFP filter set 38 HE of fluorescence microscope for AO, and a Zeiss mRFP filter set 63 HE for EB. Because of membrane integrity loss in dead cells, EB is up taken by dead cells only, but AO permeates into both dead and live cells and stains them green. Finally, dead cells are detectable in orange color in the merged images. To visualize nuclear fragmentation and condensation, cells were fixed by 70% ethanol for 20 minutes and washed with water. Then, precipitated cells were mixed with DAPI (0.1 mg/ml) in a 1:1 portion ratio to stain chromosomes. Stained cells were transferred on a glass slide and nuclei were observed under a fluorescence microscope. For microscopic analysis, a Zeiss microscope and the AxioVision 4.8 software were used to capture images, which were then analyzed using the ImageJ software.

### Nitrogen starvation and the condition in which DIM induces autophagy

The overnight culture in PMG medium (Edinburgh minimal medium (EMM) with 2% glucose and 3.75 g/L glutamate substituted for NH_4_Cl as a nitrogen source with the supplementary nucleotides and amino acids with adjusted pH ~ 6) [51] was used to get log-phase cells of the GFP-Atg8 strain. Cells from 4 ml of the culture were precipitated and washed, and medium was replaced by EMM with 1% glucose without any source of the nitrogen and incubated at 30°C. In parallel, 5 μg/ml DIM and DMSO were added to 4 ml of the overnight cultures in PMG (~0.5×10^7^ cells/ml) and incubated at 30°C in PMG. GFP-Atg8 foci were captured by a fluorescence microscope after four hours of nitrogen starvation and two hours of incubation with DIM, respectively.

## Results

### DIM reduces cell viability in log-phase cells, but not in stationary-phase cells

A high concentration of DIM (20 μg/ml) increases lifespan while a low concentration of DIM (4 μg/ml) seems to decrease lifespan in fission yeasts [44]. In humans, DIM induces apoptosis and autophagy. These facts imply that DIM may have both positive and negative effects on cell viability in fission yeast. To study the effect of DIM on fission yeast in detail, a viability assay was performed by focusing on the log-phase stage (Fig. 1A). As shown previously [44], here also, the cell viability was better when cells were incubated for 9 days in the presence of 20 μg/ml DIM. Surprisingly, we found a severe growth defect when cells were cultured for only 24 hours in the presence of DIM (Fig. 1A). DIM did not inhibit growth after additional incubation, suggesting that DIM does not inhibit growth when cells are in the stationary-phase. To test this possibility, we added DIM when the cells were in the stationary-phase. We found that DIM did not inhibit growth when we added DIM to the cells in stationary-phase (Fig. 1B). One of the explanations for the DIM resistance of the stationary-phase cells is that the cell number is high in the stationary-phase, which may result in a low effective concentration of the DIM relative to each cell. To test this possibility, we performed an acute viability assay. We adjusted cells to the same number between stationary- and log-phase cells and incubated them with DIM. To avoid cell phase-shifting after dilution of stationary-phase culture, we incubated cells for only 10 minutes with DIM. Our result showed that 10 minutes of incubation with DIM was enough to trigger cell death in log-phase cells. In contrast, the stationary-phase cells with the same cell concentration were more resistant to DIM than the log-phase cells (Fig. 1C). However, even the higher concentration (40 μg/ml) of DIM killed the stationary-phase cells, demonstrating that DIM affects viability of stationary-phase cells when the DIM concentration is very high.

### DIM induces apoptosis in log-phase cells

Based on the recent results from human studies, DIM induces apoptosis in diverse cancer cells by different mechanisms [52, 53]. We assumed that cell killing by DIM in fission yeast may be due to apoptosis induction. To know if DIM induces apoptosis in fission yeast, first, we confirmed that DIM induces cell death by staining dead and live cells (see materials and methods [27]). Dual staining by AO and EB showed that more than 80 percent of the log-phase cells were killed by DIM after 20 hrs. In contrast, dead cell abundance in treated and control cells in stationary-phase cultures did not change (Fig. 2A and 2B). These results show that DIM kills log-phase cells but not stationary-phase cells. Nuclear fragmentation is a hallmark of apoptotic cell death in fission yeasts [29]. To investigate whether DIM induces apoptosis in fission yeast, the nuclear shape was analyzed by DAPI staining. After six hours of incubation with DIM, we detected nuclear fragmentation (Fig. 2C). In contrast, we did not observe any difference in nuclei morphology in treated and control conditions in stationary-phase cells. Therefore, our results proved that DIM induces apoptosis in the log-phase cells but not in the stationary-phase cells.

**Figure 2.**
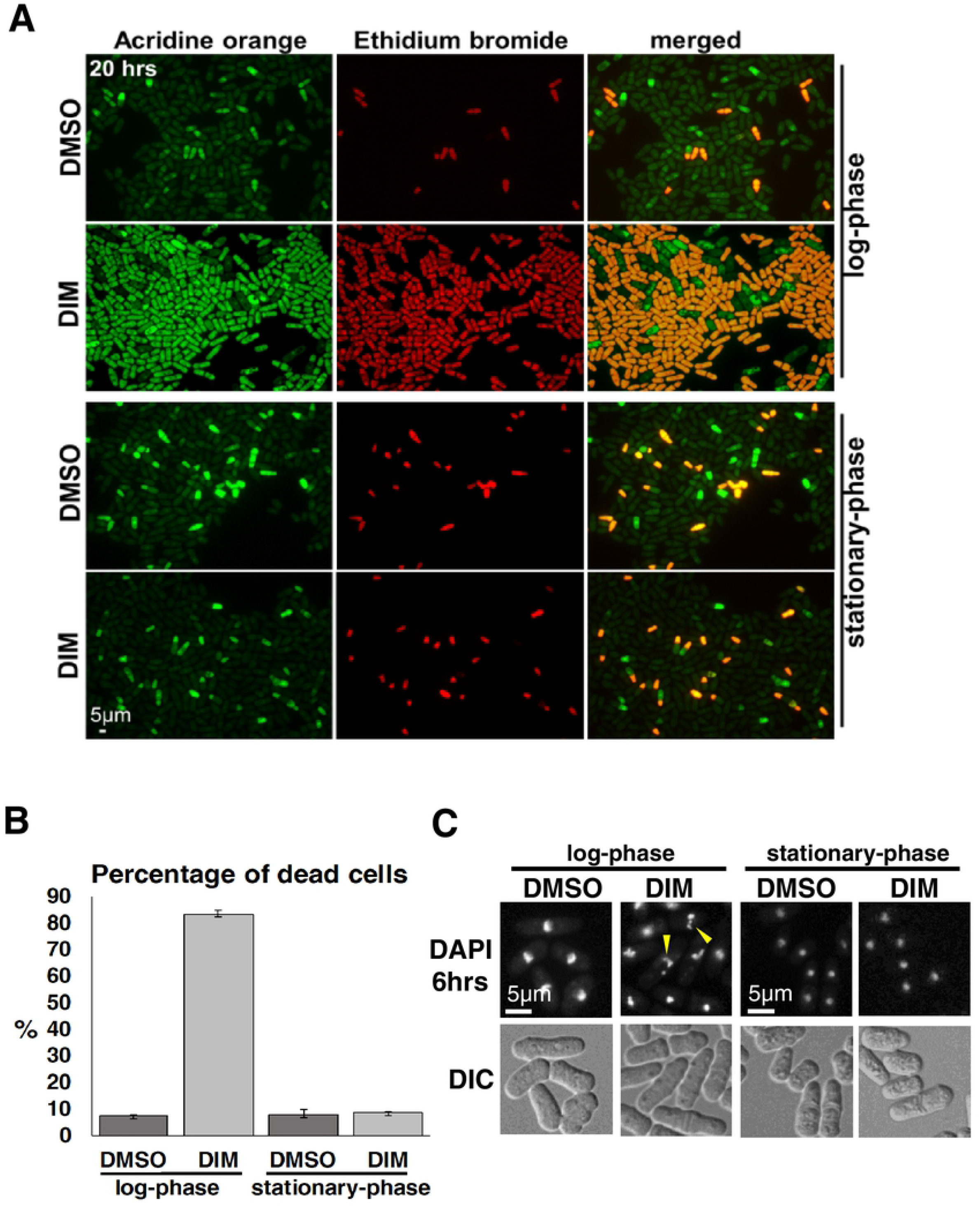
DIM induces apoptosis in fission yeast log-phase cells. **(A)** 20 hours after adding 20 μg/ml 3,3’-Diindolylmethane (DIM) to the log-phase and stationary-phase cells, the cells were stained with acridine orange (AO) and ethidium bromide (EB). Log-phase and stationary-phase cells were obtained as shown in Figure.1 A. See materials and methods for detailed condition of imaging. **(B)** The percentage of dead cells was calculated from the merged images using ImageJ software. At least 300 cells were counted with three independent experiments. **(C)** 20 μg/ml DIM was added to the log-phase and stationary-phase cells. 4,6-diamidino-2-phenylindole (DAPI) staining was used to detect the nuclear fragmentation after six hours of incubation with DIM.

### DIM induces nuclear condensation and NE disruption in log-phase cells within 10 minutes

To understand how DIM caused the reduction in viability, we analyzed nuclear morphology using cells expressing histone H2B fused to GFP (Htb1-GFP) in the presence of DIM. We observed that more than 98 percent of the log-phase cells showed nuclear condensation when the cells were incubated for 10 minutes in the presence of 20 μg/ml DIM (Fig. 3A and 3B). We also analyzed NE morphology using a strain expressing Cut11-GFP in the same condition to visualize nuclear morphology. Clearly, DIM disrupted NE in log-phase cells within ten minutes of treatment (Fig. 3C). These changes were not observed when we added DIM to the stationary-phase cells (Fig. 3D). Our results show that DIM induces nuclear condensation and NE disruption in log-phase cells, but not in stationary-phase cells within 10 minutes, which is much faster than nuclear fragmentation.

**Figure 3.**
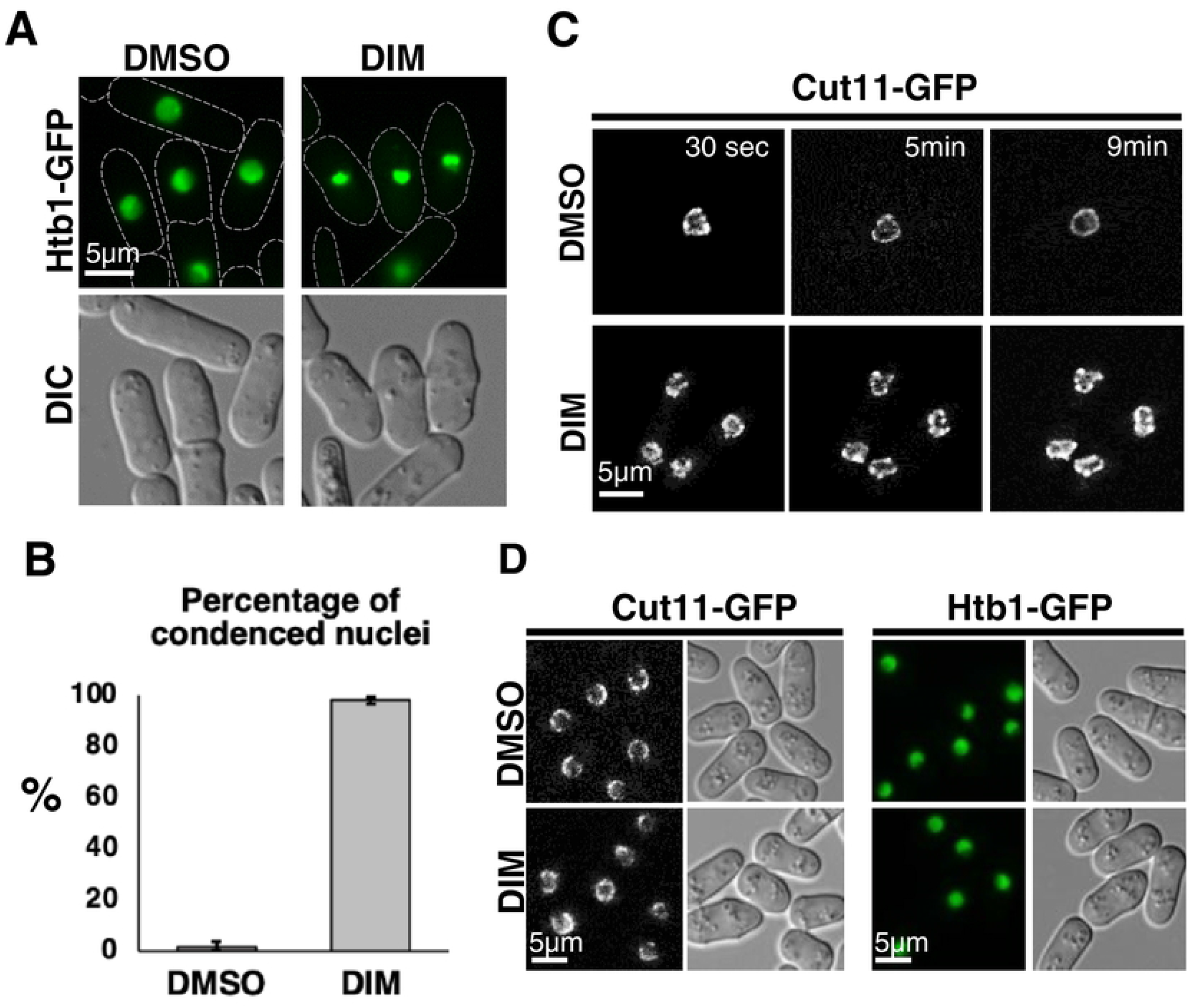
DIM induces nuclear condensation and NE disruption in log-phase cells. (**A and B)** 20 μg/ml 3,3’-Diindolylmethane (DIM) was added to log-phase cells (0.5 ~ 1 x 10^7^ cells/ml) expressing Htb1-GFP as a histone marker. Images taken after 10 min of incubation with DIM and DMSO are shown in **A**. The percentage of cells that had condensed nuclei are shown in **B**. At least 200 cells were counted with three independent experiments. **(C)** The time-lapse images of Cut11-GFP signal used as a marker for nuclear envelope (NE) are shown. Time after 20 μg/ml DIM addition to log-phase cells is shown. **(D)** 20 μg/ml DIM was added to the stationary-phase cells (10 ~ 15 x 10^7^ cells/ml) that are cultured as in Fig. 2B. After 10 min, the shape of the nuclei and NE were observed as in A and C.

### Autophagy is induced by DIM, and the autophagic pathway but not the ER stress response pathway is required for the resistance to DIM

We showed that DIM leads to apoptosis in the log-phase cells of fission yeast (Fig. 2). DIM induces autophagy in human cancer cells ([6]). To understand if DIM also induces autophagy in fission yeast, first, we used a GFP-Atg8 expressing strain. It is reported that GFP-Atg8 foci are produced by nitrogen starvation in fission yeasts [34] (Fig. 4A), which is a hallmark of autophagy induction. We observed GFP-Atg8 foci when log-phase cells were incubated with 5 μg/ml DIM for two hours (Fig. 4B). This result suggested that DIM induces autophagy in fission yeast at a low concentration (5 μg/ml). Autophagy contributes to cell viability under nitrogen starvation [25]. Next, we investigated whether autophagy contributes to survival in the presence of DIM. We used *atg7Δ* cells that have a defect in the autophagic pathway to check the viability in the presence of 5 μg/ml DIM. We found that *atg7Δ* cells are more sensitive to DIM than wild-type cells (Fig. 5A).

**Figure 4.**
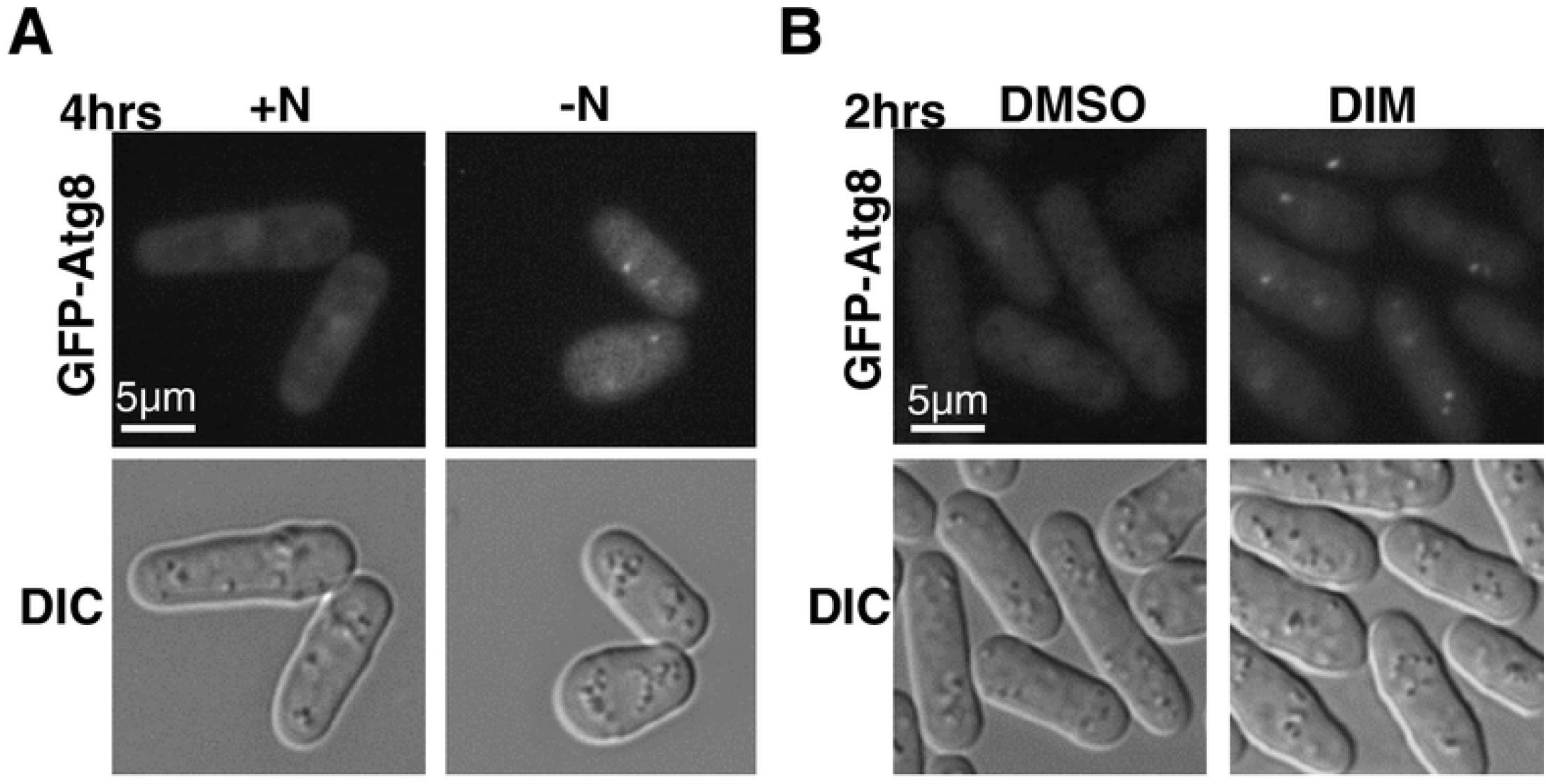
A low concentration of DIM induces autophagy in fission yeasts. **(A and B)** Cells expressing GFP-Atg8 were incubated for four hours in nitrogen starvation condition as a control **(A)** or two hours in the presence of a low concentration (5 μg/ml) of 3,3’-Diindolylmethane. **(B)** GFP-Atg8 foci were used as markers for autophagy induction. See materials and methods for detailed condition of cell cultures.

DIM also induces ER stress, which is responsible for autophagy induction in human ovarian cancer cells [6]. The next question was whether ER stress-response contributes to the viability in the presence of 5 μg/ml DIM in fission yeast. As Ire1 is the major player for ER stress response (UPR regulation) in fission yeast [28, 54], we used *ire1Δ* cells to check the DIM sensitivity. We found that *ire1Δ* cells were not sensitive to DIM, while they were sensitive to the ER stress condition induced by 2-mercaptoethanol (Fig. 5B). This data shows that ER stress response does not contribute to survival in the presence of DIM. We also investigated if ER stress response is required for autophagy induction by DIM. GFP-Atg8 foci could still be observed in the *ire1Δ* strain in the presence of DIM (Fig. 5C). These results suggested that, in contrast to the human case, ER stress response is not required for autophagy induction by a low concentration of DIM in fission yeast.

**Figure 5.**
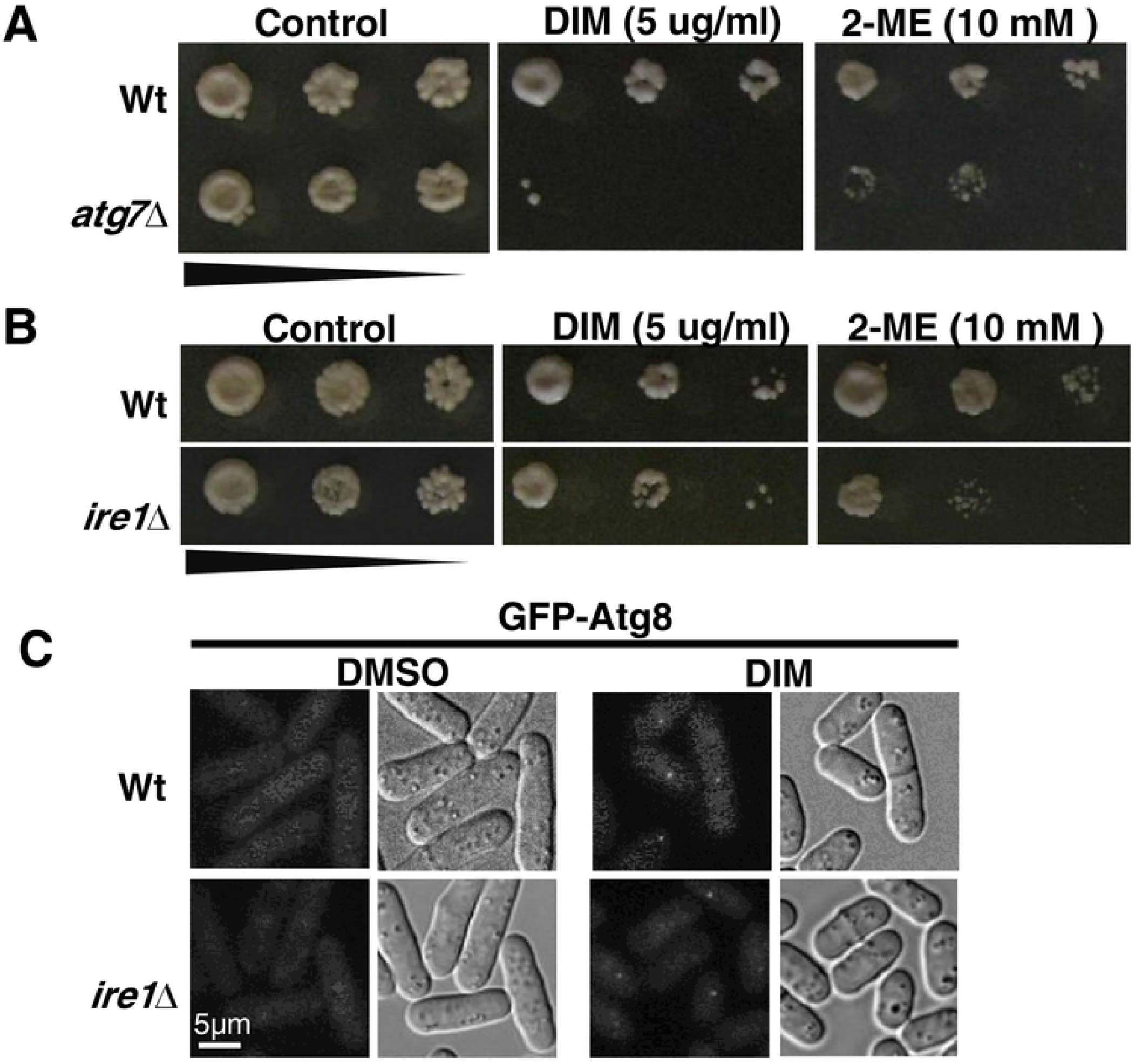
Autophagy pathway, but not ER stress response pathway, is required for the resistance to DIM. **(A and B)** Viability of log-phase cells of wild-type (Wt), *atg7Δ* **(A)** and *ire1Δ* **(B)** were studied by spot assay with five-fold serial dilutions in the presence of 5 μg/ml 3,3’-Diindolylmethane (DIM) or 10 mM 2-mercaptoethanol (2-ME). The images were taken 3-5 days after spotting. **(C)** Autophagy induction by DIM was studied using *ire1Δ* mutant expressing GFP-Atg8. The experiment was performed as shown in Figure 4. See materials and methods for details.

### Nuclear membrane protein, Lem2, is required for the resistance to DIM

We found that DIM disrupts NE rapidly (less than 10 minutes) (Fig. 3C), implying that NE could be a direct target or NE disruption might be an early event in the response to DIM. In these cases, the mutant, which has a defect in NM might be more sensitive to DIM. Lem2 is an inner NM protein and plays an important role in regulating NE membrane homeostasis [55] and chromatin anchoring to the nuclear periphery in fission yeast [38]. Therefore, we investigated whether *lem2Δ* cells are more sensitive to DIM. We found that *lem2Δ* cells are more sensitive to a low concentration of DIM (Fig. 6A). We also checked the NE morphology in the *lem2Δ* mutant in the presence of a low concentration of DIM (Fig. 6B). Unlike the high concentration of DIM (20 μg/ml), the low concentration of DIM (5 μg/ml) did not affect the NE of the wild-type strain. In contrast, NE in the *lem2Δ* mutant was significantly disrupted within four hours of incubation with DIM. These results confirmed that the NM protein, Lem2, is required for the resistance to DIM.

**Figure 6.**
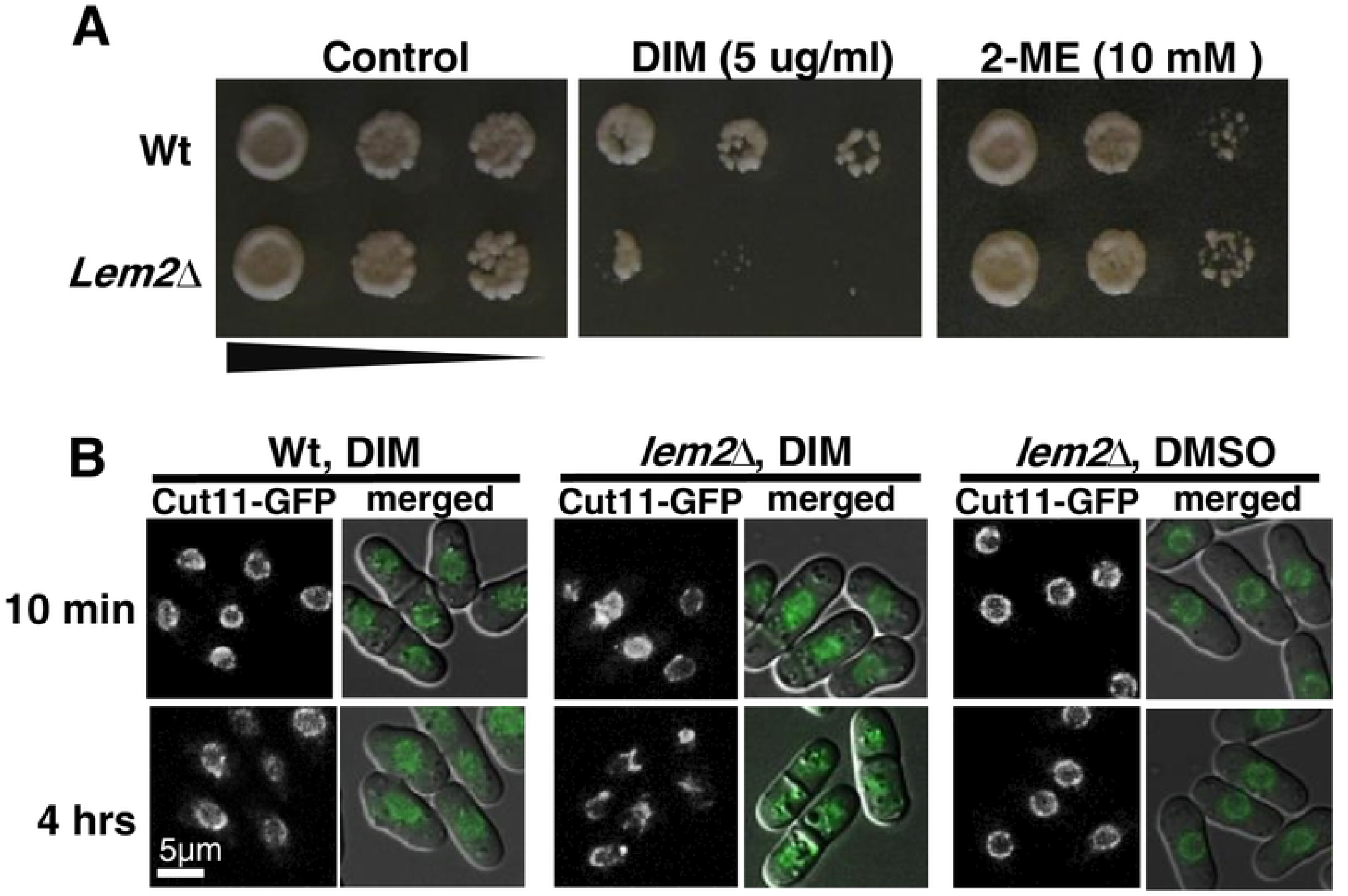
Lem2, is required for the resistance to the low concentration of DIM. **(A)** Viability of log-phase cells of wild-type and *lem2Δ* was studied by spot assay with five-fold serial dilutions in the presence of 5 μg/ml 3,3’-Diindolylmethane (DIM) or 10 mM 2-ME. Spotting assay was done as described in Figure 5A. **(B)** Fluorescent signals for Cut11-GFP protein in wild-type and *lem2Δ* log-phase cells were seen after 10 min or four hours incubation with 5 μg/ml DIM in YEA medium. Images merged with DIC are also shown.

## Discussion

### DIM induces apoptosis in fission yeasts

Recent studies in humans show that DIM is a potential anti-cancer drug that acts by the induction of apoptosis in a wide range of cancer types including the breast [11, 12, 52, 56], prostate [16], gastric [57], pancreatic [58], and hepatoma [13]. DIM also induces autophagy that substantially suppresses tumor growth and causes cancer cell death [57, 59–61]. However, the mechanisms of autophagy and apoptosis induction by DIM in humans are not fully understood.

Here we demonstrated that DIM (20 μg/ml) kills most of the log-phase cells, but not the stationary-phase cells (Fig. 2A). Additionally, we found that DIM induces nuclear fragmentation. Our results for cell killing and nuclear fragmentation strongly suggested that DIM induces apoptosis at 20 μg/ml concentration. The acute viability assay showed that 10 minutes of treatment with 20 μg/ml of DIM is enough to trigger cell killing (Fig. 1C and Fig. S1), suggesting that 10 minutes incubation with DIM is enough to induce apoptosis.

We found severe nuclear condensation and NE deformation when the log-phase cells were incubated with a high concentration of DIM for 10 minutes (Fig. 3A and B). We did not see nuclear fragmentation, nuclear condensation, or NE disruption, when we decreased the drug concentration to 5 μg/ml (Fig. S2). These results show that DIM effects are dose-dependent in log-phase cells. We speculate that DIM may inhibit some of the membrane proteins that are required for NE integrity, or inhibit control of the protein transport between nucleus and cytoplasm at the 20 μg/ml concentration. Therefore, by disrupting the NE integrity, DIM may cause the mis-localization of the nuclear or cytoplasmic protein. Subsequently, this mis-localization may change nuclear homeostasis and induce nuclear condensation. Nuclear condensation and NE disruption are the signs of apoptosis induction in fission yeast [28, 29, 62]. However, in our experiments, nuclear condensation and NE disruption happened much earlier than previous observations [28, 29, 62]. The disruption of nuclear transport and disassembly of the nuclear pore complex (NPC) occurs before caspase-9 activation in humans [63]. Therefore, it could be possible that the nuclear condensation and NE disruption by DIM in fission yeast, in the first 10 minutes of incubation, happen before apoptosis is induced. NE disruption by DIM and possible subsequent defect in protein transport between the nucleus and the cytoplasm might be a reason for apoptosis induction. However, it remains unclear how DIM induces apoptosis in fission yeast and further studies are necessary to understand the mechanism of apoptosis induction by DIM.

### Mechanism of autophagy induction by DIM

Similar to the results in human studies [6, 57], we found that DIM induces autophagy in fission yeast (Fig. 4B) when we used the low concentration of DIM (5 μg/ml). Based on our data, the autophagy pathway contributes to the resistance to a low concentration of DIM (Fig. 5A). Nitrogen starvation [32, 34], sulfur depletion [35], or ER stress [32] induce autophagy in fission yeast. It remains unclear how DIM induces autophagy in fission yeast. However, we found the autophagy induction by DIM is Ire1-independent, demonstrating that autophagy induced by DIM is not likely ER stress response-dependent. The *ire1Δ* strain did not show DIM sensitivity while it was ER stress-sensitive (Fig. 5B). These results suggest that ER stress response is not required for the cell viability in the presence of a low concentration of DIM. DIM might mimic nitrogen starvation or sulfur depletion or alternatively, DIM might induce autophagy by an unidentified pathway.

### Lem2 is required for NE integrity in the low concentration of DIM

Finally, we showed that the inner NM protein, Lem2, is critical for cellular viability at a DIM concentration that induces autophagy (Fig. 6A). The concentration of DIM that induced autophagy resulted in NE disruption in *lem2Δ* cells but not in wild-type cells (Fig. 6B). Lem2 acts as a barrier to membrane flow between the NE and other parts of the cellular membrane system [40]. On the other hand, the amount of C24:0 fatty acid, which is important for the survival of yeast cells [64], is reduced in the absence of both Lem2 and Bqt4 in fission yeast [65]. Therefore, the physical property of the NM including fatty acid composition and possibly membrane protein composition may be affected in *lem2Δ* cells. This could be one of the possible reasons why NE in *lem2Δ* cells is more susceptible to the lower concentration of DIM. Also, *lem2Δ* cells have defects in the recruitment of proteins such as Vps4 and Cmp7 to NE and in repairing the holes of NE by ESCRT-III [39]. Therefore, DIM may make holes in the NE of wild-type cells that could be repaired. In contrast, in the *lem2Δ* strain with defects in Vps4 expression and Cmp7 localization, ESCRT-III may be unable to seal the NE. This could be another reason for the DIM sensitivity in *lem2Δ* cells.

### Stationary-phase cells are resistant to DIM

Our result showed that there is a difference between log-phase and stationary-phase cells in response to DIM (Fig. 1). We found that the log-phase cells, but not stationary-phase cells, are sensitive to the higher concentration of DIM (20 μg/ml). The ratio of drug molecules per cell is not the reason for the difference in viability of log-phase and stationary-phase cells in response to DIM (Fig. 1C). Also, NE, nuclear morphology, and cell survival were not affected at the higher concentration of DIM (20 μg/ml) in stationary-phase cells (Fig. 2B and Fig. 3).

Our results suggest that NE could be one of the early targets for DIM in log-phase-cells (Fig. 3C, 6B). Therefore, the difference in the nature of NE and protein composition in the NE between log-phase and stationary-phase cells could be the reason for the difference in sensitivity. The NE protein Ish1 has a higher level of expression in stationary-phase than in log-phase [66]. Therefore, Ish1 may be a candidate for the resistance to DIM in stationary-phase cells. However, it is possible that DIM targets the unknown NE-independent protein(s) in log-phase cells, which may be absent in stationary-phase cells.

In conclusion, we showed for the first time that DIM induces apoptosis and autophagy in fission yeast, which was previously reported in humans. The mechanism by which DIM induces apoptosis and autophagy may be conserved in yeast and humans, which helps to study it easily in a unicellular organism such as the fission yeast. Also, the difference between apoptosis and autophagy induction due to DIM concentration implies the importance of the dose-dependent manner in which DIM affects the final killing or survival of the cells. Also, we showed that NE could be one of the early targets of DIM. The understanding of apoptosis and autophagy mechanism by DIM in fission yeast may be helpful for human cancer and longevity research. We think that study of the NE structure could be a good starting point for these cases.

## Acknowledgment

We are grateful to Dr. Mitsuhiro Yanagida, Dr. Atsushi Matsuda, Dr. Yasushi Hiraoka, Dr. Peter Walter, Dr. Ayumu Yamamoto, Dr. Yoshinori Watanabe, and Dr. Takeshi Sakuno, Dr. A. Francis Stewart, and Sayaka Suzuki for the strains and plasmids, respectively. We are also thankful to Dr. Hizlan Hincal Agus for his helpful guidance.

